# A feed-forward relay between Bicoid and Orthodenticle regulates the timing of embryonic patterning in *Drosophila*

**DOI:** 10.1101/198036

**Authors:** Rhea R. Datta, Jia Ling, Jesse Kurland, Xiaotong Ren, Zhe Xu, Gozde Yucel, Jackie Moore, Leila Shokri, Isabel Baker, Timothy Bishop, Paolo Struffi, Rimma Levina, Martha L. Bulyk, Robert J. Johnston, Stephen Small

## Abstract

The K50 homeodomain (K50HD) protein Orthodenticle (Otd) is critical for anterior patterning and brain and eye development in most metazoans. In *Drosophila melanogaster*, another K50HD protein, Bicoid (Bcd), has evolved to replace Otd’s ancestral function in embryo patterning. Bcd is distributed as a long-range maternal gradient and activates transcription of a large number of target genes including *otd*. Otd and Bcd bind similar DNA sequences *in vitro*, but how their transcriptional activities are integrated to pattern anterior regions of the embryo is unknown. Here we define three major classes of enhancers that are differentially sensitive to binding and transcriptional activation by Bcd and Otd. Class 1 enhancers are initially activated by Bcd, and activation is transferred to Otd via a feed-forward relay (FFR) that involves sequential binding of the two proteins to the same DNA motif. Class 2 enhancers are activated by Bcd, and maintained by an Otd-independent mechanism. Class 3 enhancers are never bound by Bcd, but Otd binds and activates them in a second wave of zygotic transcription. The specific activities of enhancers in each class are mediated by DNA motif variants preferentially bound by Bcd or Otd, and the presence or absence of sites for cofactors that interact with these proteins. Our results define specific patterning roles for Bcd and Otd, and provide mechanisms for coordinating the precise timing of gene expression patterns during embryonic development.

## Introduction

Animal body plans are established in large part by transcriptional networks that provide positional information for specific cell fates during embryogenesis [1]. The evolution of body plans is driven by alterations to these networks, including changes to *cis*-regulatory elements and neofunctionalization of transcription factors (TFs) after gene duplication [2-6]. At the transcriptional level, most network interactions involve direct binding of TFs to binding sites in the enhancers of target genes. Individual binding events are then integrated into spatial and temporal patterns of expression that create positional information along the major axes during development. Understanding the mechanisms controlling the dynamics of gene expression remains a challenge, in part because TFs exist in families of closely related proteins that are coexpressed and bind very similar sequence motifs *in vitro* [7]. While possible mechanisms by which TF specificity is achieved (TF-preferred binding sequences, cofactor binding, etc.) have been proposed [8], it is unclear how binding events are coordinated *in vivo* so that each protein can activate its own specific gene targets.

The early *Drosophila melanogaster* embryo provides a unique system to study the spatiotemporal complexity of transcription factor-DNA interactions. *Drosophila* and other Cyclorrhaphan Diptera develop via a “long germ” mode, in which all segments of the body plan are specified and positioned during the syncytial blastoderm stage [9-11] In the fruit fly, the maternal transcription factor Bicoid (Bcd) sets up the first positional instructions for anterior patterning [12]. *bcd* mRNA is sequestered at the anterior pole of the mature oocyte via sequences in its 3’ UTR [13]. After fertilization, translation and diffusion establish an anterior gradient of Bcd protein [14, 15] (Fig 1A-B). Bcd contains a homeodomain (HD), and is a transcriptional activator of more than 50 target genes [16-20], which form a network of downstream regulatory interactions that precisely pattern the embryo. Embryos lacking Bcd fail to form any anterior structures, including all cephalic and thoracic segments [21]. They also show duplications of posterior structures (ex: Filzkoerper) in anterior regions [21]. While the majority of its targets are expressed during and after cellularization, the Bcd protein gradient is only active during the syncytial blastoderm stage, prior to cellularization.

**Figure 1.**
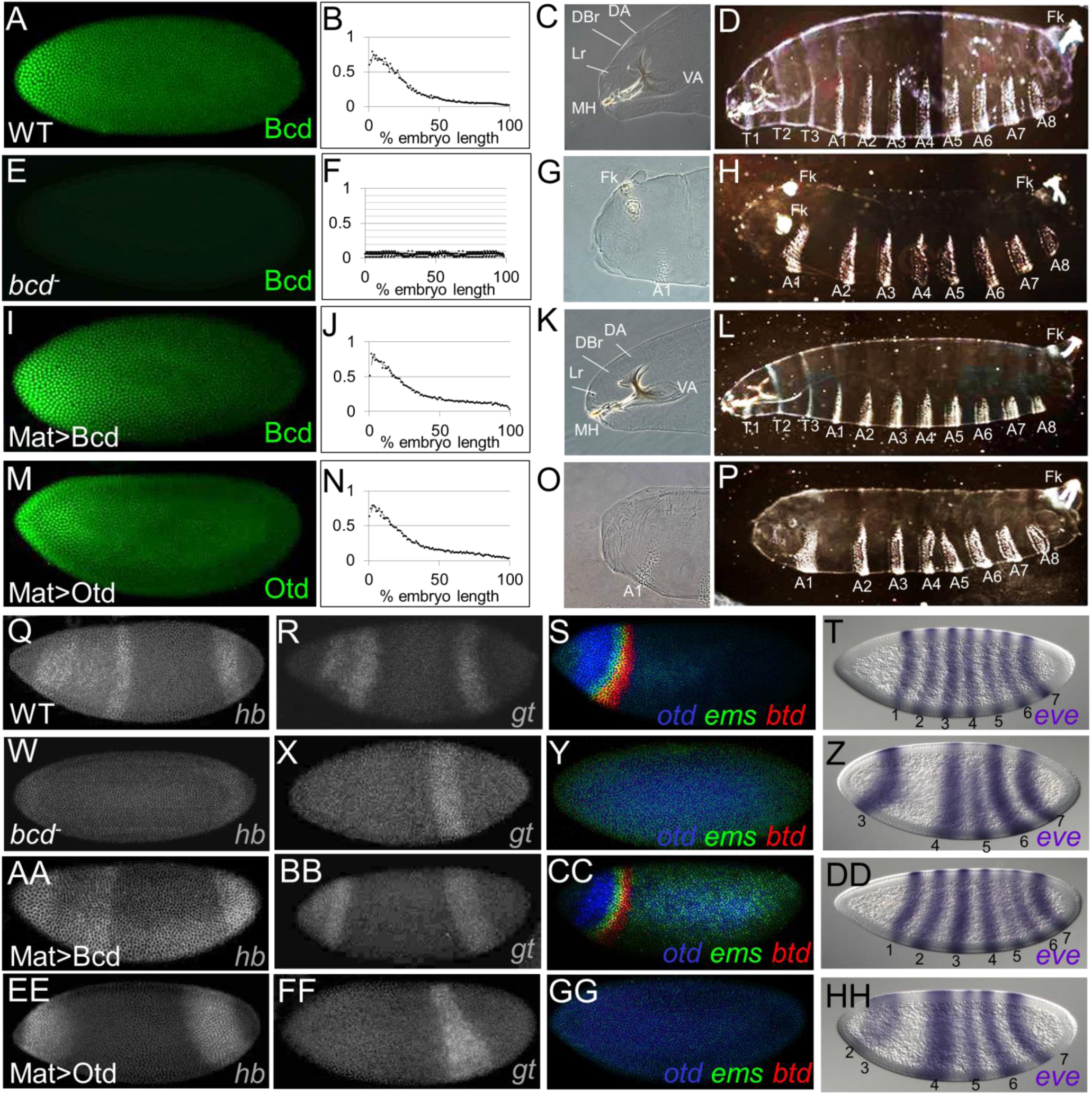
A maternal Otd gradient (Mat>Otd) cannot replace most Bcd-like functions. **(A-P)** Protein expression patterns (A, E, I, M), gradient quantifications (averaged from five embryos; B, F, J, N), anterior cuticle structures (C, G, K, O), and whole larva cuticles (D, H, L,P) are shown for wild-type (A-D), *bcd* mutants (E-H), *bcd* mutants containing the Mat>Bcd transgene (I-L), and *bcd* mutants containing the Mat>Otd transgene (M-P). Labeled structures in anterior regions (C, G, K, O) include the dorsal arm (DA), dorsal bridge (DBr), labrum (Lr), mouthhooks (MH), ventral arm (VA), Filzkoerper (Fk), and the 1^st^ abdominal segment (A1). (D, H, L, P) Thoracic (T1-3) and abdominal (A1-8) segments are labeled. **(Q-HH)** mRNA expression patterns for six Bcd target genes (*hb*, *gt*, *otd*, *ems*, *btd*, and *eve*) in wild-type (Q-T), *bcd* mutants (W-Z), *bcd* mutants containing the Mat>Bcd transgene (AA-DD), and *bcd* mutants containing the Mat>Otd transgene (EE-HH). Assayed target genes are labeled in the lower right hand corner of each panel. (T, Z, DD, HH) Numbers correspond to eve stripes. All embryos in this paper are oriented with anterior to the left.

Despite its critical functions in *Drosophila*, Bcd is not well conserved, even within insects. Rather, Bcd arose after a recent gene duplication event, and rapidly evolved to play an important role in embryonic patterning [22, 23]. In three species lacking Bcd, *Tribolium castaneum* (*Tc*), *Nasonia vitripennis* (*Nv*) and *Acyrthosiphon pisum* (*Ap*), at least some of the anterior patterning functions of Bcd are performed by Orthodenticle (Otd) [24-26]. *otd* is maternally expressed in these species, and its disruption causes severe *bcd*-like defects in anterior patterning. In *Drosophila*, *otd* has evolved to become a zygotic target gene of Bcd-dependent activation [18]. Loss of *Drosophila otd* causes embryonic lethality, but *otd* mutants show cephalic defects that are much less dramatic than the complete loss of anterior structures observed in *bcd* mutants [18, 27]. Later in development Otd is critical for central nervous system and eye development [27], roles that are conserved in vertebrates including humans [28].

Otd and Bcd each contain a lysine at amino acid position 50 (K50) of their respective homeodomains, and bind *in vitro* to the consensus sequence TAATCC [29, 30]. The K50 residue and preference for TAATCC are conserved among all Otd homologs [28], and is thus ancient, but the ancestral protein that gave rise to Bcd was a Hox3 protein with a glutamine (Q) at HD position 50 (Q50) [23]. This suggests that an important step in Bcd’s evolution was the conversion of Q50 in the ancestor to K50, which changed its DNA-binding preference and allowed it to usurp some of the anterior patterning roles played by Otd in ancestral insects. All known Bcd target gene enhancers contain multiple copies of TAATCC [16]. We hypothesize that Otd regulated many of these enhancers in the ancestral species that gave rise to *Drosophila*, and that it regulates a similar battery of enhancers in extant insect species that do not contain Bcd. If this is the case, perhaps Otd can replace many of Bcd’s functions in *Drosophila* if maternally expressed and distributed in an anterior gradient.

Here we test whether Otd can provide Bcd-like patterning functions through a transgenic gene replacement assay. We also present a comprehensive comparison of the *in vitro* binding preferences and *in vivo* binding profiles of Bcd and Otd. We demonstrate that the two proteins bind sequentially to enhancers in feed-forward relays (FFRs), in which Bcd-binding initiates target gene activation, and Otd-binding maintains expression after the maternal Bcd gradient decays. Each protein also binds independently to distinct enhancers. We present evidence that Otd- and Bcd-specific binding activities *in vivo* are controlled by two mechanisms: subtle differences in binding preference and protein-specific interactions with cofactors. Our results define specific roles for Bcd and Otd in embryonic patterning in *Drosophila*, and shed light on the molecular mechanisms that alter regulatory networks during evolution.

## Results

### A maternal gradient of Otd cannot replace Bcd in *Drosophila*

To assess the functional similarity between Bcd and Otd, we tested whether Otd could mediate Bcd-like activities using a gene replacement assay (Fig. 1). Coding regions for both Bcd and Otd were each inserted into transgenes containing the *bcd* promoter (for maternal expression) and the *bcd* 3’ UTR (for anterior mRNA localization), and were integrated into the same genomic position [31]. These constructs (designated Mat>Bcd and Mat>Otd) were crossed into *bcd*^*E1*^ null mutant females, and embryos laid by those females were assayed for RNA and protein expression. mRNAs from the transgenes were maternally expressed and localized to the anterior pole region (Supp. Fig. 1A). Antibody stains showed very similar gradients of Bcd and Otd in early embryos containing the transgenes, and the Bcd gradient from the rescue transgene was indistinguishable from the endogenous (WT) gradient (Fig. 1A, B, I, J, M, N).

Wild-type embryos develop cephalic structures and three thoracic segments in anterior regions during embryogenesis (Fig.1C, D). All these structures are missing in embryos laid by *bcd* mutant females, and posterior structures [Filzkoerper (Fz)] are duplicated near the anterior pole. (Fig. 1G, H). At the molecular level, *bcd* mutants fail to activate all Bcd-target genes, including *hunchback* (*hb*), *giant* (*gt*), *otd*, *buttonhead*, (*btd*), *empty spiracles* (*ems*), and *even-skipped* (*eve*) stripes 1 and 2. (Fig 1W-Z, compare to 1Q-T), and fail to repress translation of Cad in anterior regions (Supp. Fig. 1B).

As expected, when the control construct (Mat>Bcd) was crossed into embryos lacking Bcd, it completely rescued all morphological defects (Fig. IK, L), and 95% of the embryos developed into fertile adults. This construct also activated the expression patterns of all six tested Bcd target genes (Fig. 1AA-DD). In contrast, when the Mat>Otd construct was crossed into *bcd* mutants, none of the morphological structures missing in *bcd* mutants structures were rescued (Fig. 1O, P). However, we did detect the suppression of ectopic posterior Filzkoerper (Fig. 1O; compare to 1G). Mat>Otd activated only two Bcd target genes [*hunchback* (*hb*) and *even-skipped stripe-2* (*eve-2*)], but failed to activate four others [*giant* (*gt*), *otd*, *buttonhead*, (*btd*), and *empty spiracles* (*ems*)] (Figure 2EE-HH), and failed to translationally repress *caudaI* (Supp. Fig. 1B). We also tested whether Mat>Bcd and Mat>Otd could activate 24 other Bcd-dependent reporter genes [16]. The Mat>Bcd construct activated expression of all 24 of these reporters, while Mat>Otd activated only two (Supp. Fig. 2). Taken together, these results show that Otd cannot rescue most Bcd functions when provided maternally, despite the fact that the two proteins bind very similar DNA sequences *in vitro*.

**Figure 2.**
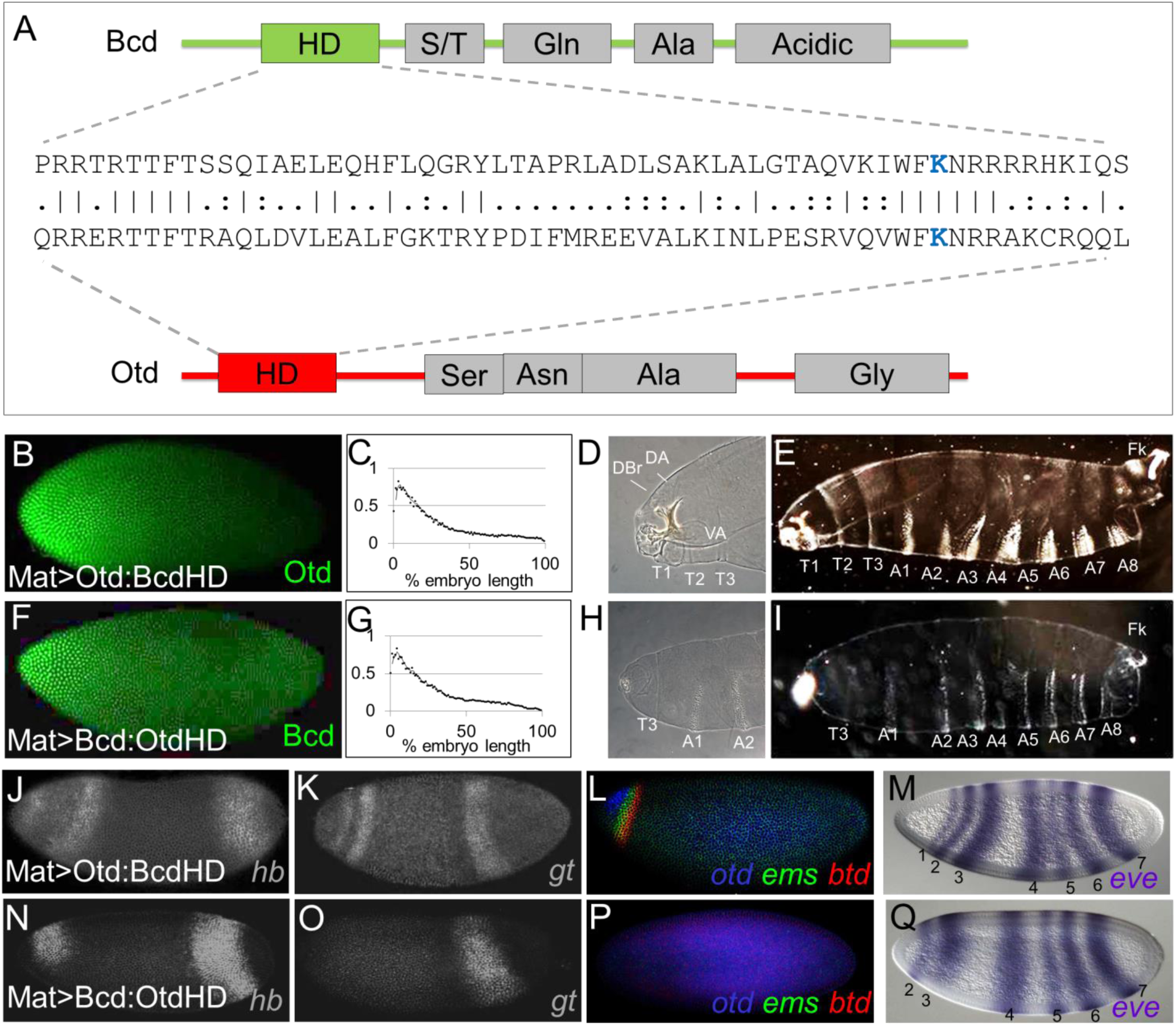
**(A) A structural comparison of the Bcd and Otd proteins.** Schematic representations show positions of the homeodomains (HDs) and mapped activation domains (gray boxes). An amino acid sequence comparison of the HDs is shown in the middle. Identical amino acids are indicated by vertical lines, and similarities are shown as two dots. The critical lysine at position 50 (K50) is shown in blue. **(B-I) Testing HD swap chimeras for Bcd-like activities. (B-I)** Protein expression patterns (B, F), gradient quantifications (averaged from five embryos; C, G), anterior cuticle structures (D, E, H, I), and whole larva cuticles (E, I) are shown for *bcd* mutants containing the Mat>Otd:BcdHD transgene (B-E), and *bcd* mutants containing the Mat>Bcd:OtdHD transgene (F-I). Labeled structures in anterior regions (D, H) include the dorsal arm (DA), dorsal bridge (DBr), ventral arm (VA), thoracic segments (T1-T3), and the 2 abdominal segments (A1 and A2). (E, I) Thoracic (T1-3) and abdominal (A1-8) segments are labeled. **(J-Q)** mRNA expression patterns for six Bcd target genes (*hb*, *gt*, *otd*, *ems*, *btd*, and *eve*) in *bcd* mutants containing the Mat>Otd:BcdHD transgene (J-M), and *bcd* mutants containing the Mat>Bcd:OtdHD transgene (N-Q). (M, Q) Numbers correspond to eve stripes. Assayed target genes are labeled in the lower right hand corner of each panel.

### HDs mediate distinct *in vivo* activities of Otd and Bcd

The inability of Mat>Otd to activate most Bcd target genes or repress Cad is consistent with its failure to rescue anterior segments in embryos lacking Bcd. This result is not surprising in view of the fact that the evolved Bcd and Otd coding regions show very little sequence conservation (38% sequence identity within their homoeodomains (HDs) (Fig. 2A), and no detectable homology outside the HD). To map regions of Bcd and Otd that mediate their distinct functions, we generated rescue constructs with chimeric proteins in which the DNA-binding homeodomains (HDs) were precisely swapped (Mat>Bcd:OtdHD and Mat>Otd:BcdHD, Fig. 2B, C, F, G). If the structural differences preventing Otd from rescuing *bcd* mutants lie inside its HD, inserting the Bcd HD into the Otd protein (Mat>Otd:BcdHD) should cause a strong rescue of the phenotype. If those differences lie outside the HD inserting the Otd HD into the Bcd protein (Mat>Bcd:OtdHD) should result in a strong rescue.

Inserting the Bcd HD into the Otd protein (Mat>Otd:BcdHD) caused a dramatic rescue of anterior structures (Fig. 2D, E). Although rescue was incomplete (embryos died before hatching), all embryos containing the Mat>Otd:BcdHD formed three thoracic segments, and at least some identifiable cephalic structures (Fig. 2D, E). Mat>Otd:BcdHD also activated all six tested Bcd target genes (Fig. 2J-M), and repressed Cad translation (Supp. Fig. 1B). In contrast, the reciprocal swap (Mat>Bcd:OtdHD) caused very little rescue of the morphological defects of *bcd* mutants (Fig. 2H, I), and activated only *hb* and *eve2*, similar to rescue by Mat>Otd (Fig. 2N-Q compare to Fig.1 EE-HH).Together, these results indicate that important functional differences between the two proteins lie within their HDs, and regions outside their HDs are largely interchangeable.

### *In vivo* and *in vitro* binding activities of Bcd and Otd

The result that the Bcd and Otd HDs are not interchangeable seems in conflict with the observation that Bcd and Otd bind the same TAATCC consensus sequence *in vitro* [7]. However, it is possible that preferential binding to suboptimal sites might enable their distinct functionalities *in vivo*. Alternatively, the Bcd and Otd HDs might differentially interact with cofactors. To compare the binding activities of Bcd and Otd in wild type embryos, we performed ChIP-Seq experiments. To determine the best time intervals for embryo collection, we double-stained embryos at five consecutive time-points after egg laying (AEL) using anti-Bcd and anti-Otd antibodies in wild type embryos. These experiments show the Bcd gradient at Stage 4 (S4), but no Otd expression at this stage (Fig. 3A). Otd protein is first visible at early Stage 5 (S5, Fig. 3B) and its expression increases at mid- to late-S5, when both Bcd and Otd protein are strongly expressed (Fig. 3C). After cellularization during stages 6-8 (S6-8), Bcd protein becomes undetectable (Fig. 3D), while Otd expression is maintained in the head and up-regulated along the ventral midline (Fig. 3D-E’)

**Figure 3.**
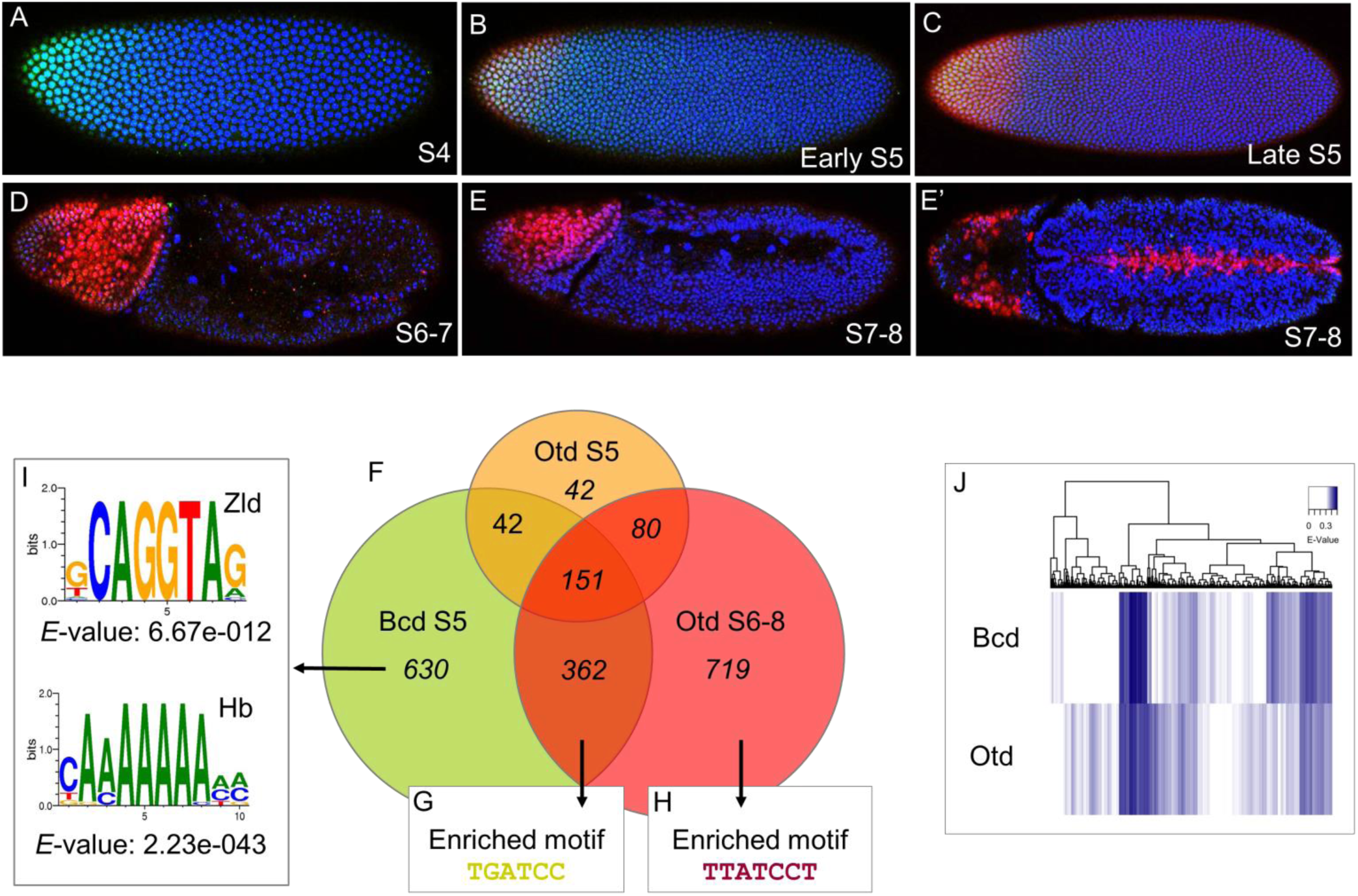
**(A-E’) A temporal comparison of Bcd and Otd expression.** All embryos are stained for Bcd protein (green), Otd protein (red), and DAPI (blue), which marks individual nuclei (blue). The embryos shown represent a temporal series from A (youngest) to E (oldest). The ages of embryos are labeled by stage (S). **(F-I) Genome-wide binding activities of Bcd and Otd.** (F) A Venn diagram showing the number Bcd and Otd peaks in euchromatin (FDR<5%) in wild type embryos at the S5 and S6-8 time-points. Motifs enriched in each dataset are indicated (G-I). (J) **Heat map comparisons of 9-mers bound by Bcd and Otd** Hierarchical clustering analysis of E-scores. All 9-mers bound by Bcd and Otd with an E-value>0 are represented. Every position on the heat-map is a single 9-mer.

Based on these experiments we performed ChIP-Seq on collections of embryos at S5 when the two proteins are co-expressed, and at S6-8, when only Otd is detectable (see Methods). We detected 1185 Bcd-bound peaks in S5 embryos (Fig. 3F). 99% of these peaks mapped to euchromatic regions of the genome, and 65 of 66 previously known Bcd-dependent enhancers [16] were detected in this experiment. Otd bound to 524 peaks at this time-point, but only 60% (315 peaks) mapped to euchromatin. The remaining 40% (209 peaks) mapped to heterochromatic or uncharacterized regions of the genome. The Otd early euchromatic binding included regions that overlapped with Bcd-binding peaks. Only 42 peaks (13% of all Otd early peaks) bound by Otd at S5 mapped to euchromatic regions. At the later time-point, no significant Bcd-binding was detected, but Otd bound to 1312 peaks, 98% of which mapped to euchromatic regions (Fig. 3F).

Comparisons of the ChIP-Seq profiles at both time-points showed that Bcd and Otd bind to 630 and 719 unique peaks respectively (Fig. 3F), supporting the observation that the two proteins are not functionally interchangeable *in vivo* (Fig. 1). The molecular basis for differential binding *in vivo* is not clear, but one possibility is that the Bcd and Otd HDs have inherent binding preferences that were not detected in previous in vitro binding studies. To test this, we used universal protein-binding microarrays (PBMs), in which purified GST-tagged Otd and Bcd HDs were tested for binding to all possible 9-mer nucleotide sequences ([32]; Methods). Previous work has shown that if a *Drosophila* HD is bound to a 9-mer sequence in a PBM with an E-score>0.31 it is likely to be a functionally relevant site *in vivo* [33]. A comparative heat-map of Bcd and Otd binding to every possible 9-mer is shown in Fig. 4J. Only 9-mers with an E-score>0.31 are shown in color. The PBM E-score binding profiles indicate differences in binding preferences between Bcd and Otd (Fig. 4J).

**Figure 4.**
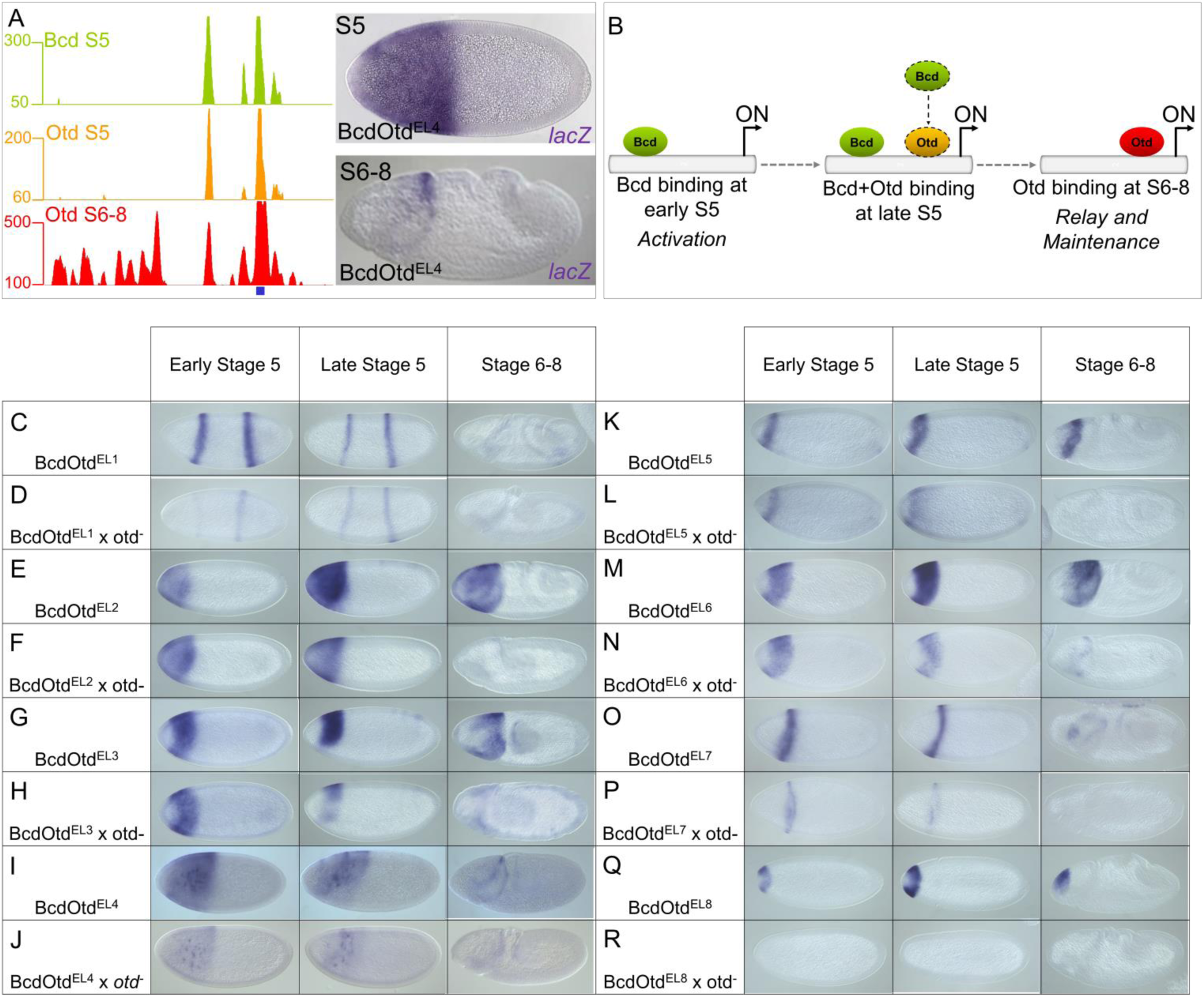
Temporal regulation of feed-forward relay enhancer activity by Bcd and Otd. **(A)** ChIP-seq peaks for Bcd and Otd around the *hb* locus. The position of a known Bcd-dependent enhancer is shown as a blue box. This enhancer drives expression early at S5 and later at S6-8. **(B)** A model for a feed-forward relay coordinated by Bcd and Otd. **(C-T) Testing feed-forward relay enhancers** in *otd* mutant embryos. Reporter gene expression patterns in wild-type embryos (C, E, G, I, K, M, O, Q) and *otd* mutant embryos (D, F, H, J, L, N, P, R). Embryos are shown at 3 time-points, as indicated on top.

The CHIP-Seq experiments also detected 513 peaks that were bound by Bcd early and then by Otd at the late time-point (Fig. 3F). These overlapping peaks include 53 of 66 previously characterized Bcd-dependent enhancers ([16]; Fig. 4A, Supp. Fig. 3). We hypothesize that these enhancers are regulated by a common feed-forward relay (FFR) that integrates the activities of Otd and Bcd. In this model genes controlled by these relay enhancers are initially activated by Bcd, and then regulated by Otd after the Bcd gradient decays (Fig. 4B), and the transfer of control from Bcd to Otd would effectively extend the time of expression for a specific set of target genes. We classify these enhancers as BcdOtd^EL^ where E= bound **e**arly (S5) and L= bound **l**ate (S6-8), and use a consistent nomenclature throughout the rest of the paper.

### A feed-forward relay regulates enhancers bound by Bcd and Otd

To test the feed-forward relay hypothesis, we examined the activities of 8 candidate relay enhancers that were bound by both Bcd and Otd, expressed in anterior regions, and continuously active during the period between S5 and S6-8 (Fig. 4B-R, Supp. Fig. 2). In a previous study, we showed that removing Bcd function (which also removes Otd) completely abolished expression driven by all these enhancers [16]. If Otd is involved in maintaining the expression patterns driven by these enhancers, removing its function should cause a loss or reduction of expression at S6-8. To ablate Otd function, we used CRISPR-Cas9-mutagenesis to delete the Otd HD in the endogenous locus (Methods). This mutation caused embryonic lethality, but did not affect the shape of the Bcd gradient (Supp. Fig. 4).

Expression patterns driven by the eight candidate relay enhancers were assayed in *otd* mutant embryos at three time-points, early S5 (high Bcd, low Otd), late S5 (high Bcd, high Otd), and S6-8 (no Bcd, high Otd). For 6 of the 8 tested enhancers, removing Otd caused a substantial reduction in expression at late NC14 and/or a complete loss of expression at S6-8, but had little effect on early activation (Fig. 4C-H, K-P). One of these enhancers was the EHE enhancer from the *otd* gene itself (BcdOtd^EL6^, Supp. Fig. 3), and the strong reduction of expression driven by that enhancer (Fig. 4M, N) suggests that maintenance of the normal *otd* pattern is mediated by a positive autoregulatory loop. The other two reporter genes (BcdOtd^EL4^ and BcdOtd^EL8^) tested in *otd* mutants showed strong reductions even in early NC 14 (Fig. 4I, J, Q, R, Supp. Fig. 3), suggesting that Otd is required for the initial activation of some Bcd target genes. We define these as “relay enhancers” - they are bound and activated by Bcd prior to S5 and are then maintained by Otd in later development. Taken together, these results and the large number of shared peaks in the ChIP-Seq experiments suggest that the feed-forward relay (FFR) between Bcd and Otd is a common mechanism for extending the timing of expression of a large subset of target genes in anterior regions of the embryo. In this relay mechanism, Otd’s primary role is the maintenance of gene expression.

### The Bcd-Otd FFR involves sequential binding to a suboptimal site

One possibility is that FFR-regulated enhancers contain specific sequence motifs that facilitate sequential binding of Bcd and Otd. Using a discriminative motif search (see Methods), we identified a single base variant of the K50 consensus, TGATCC, which is enriched in peaks bound by both proteins compared to peaks bound by Bcd or Otd alone (Fig. 3G). This site does not contain the TAAT core recognition site preferred by most HD-containing transcription factors [7, 30]. This TGATCC motif appears in 62% of the overlapping peaks compared to 23% of Bcd S5 only peaks (p=0) and 38% of Otd S6-8 only peaks (p=9.05E-08). We also searched the data from our PBM experiments to see how this single base change affects the binding preferences of Bcd and Otd (Fig. 5). This search showed that TGATCC-containing 8-mers are suboptimal (less strongly bound by both proteins) compared to the TAATCC consensus (Fig. 5A). It also appears that Bcd prefers TGATCC-containing 8-mers more strongly than does Otd (Fig. 5B).

**Figure 5.**
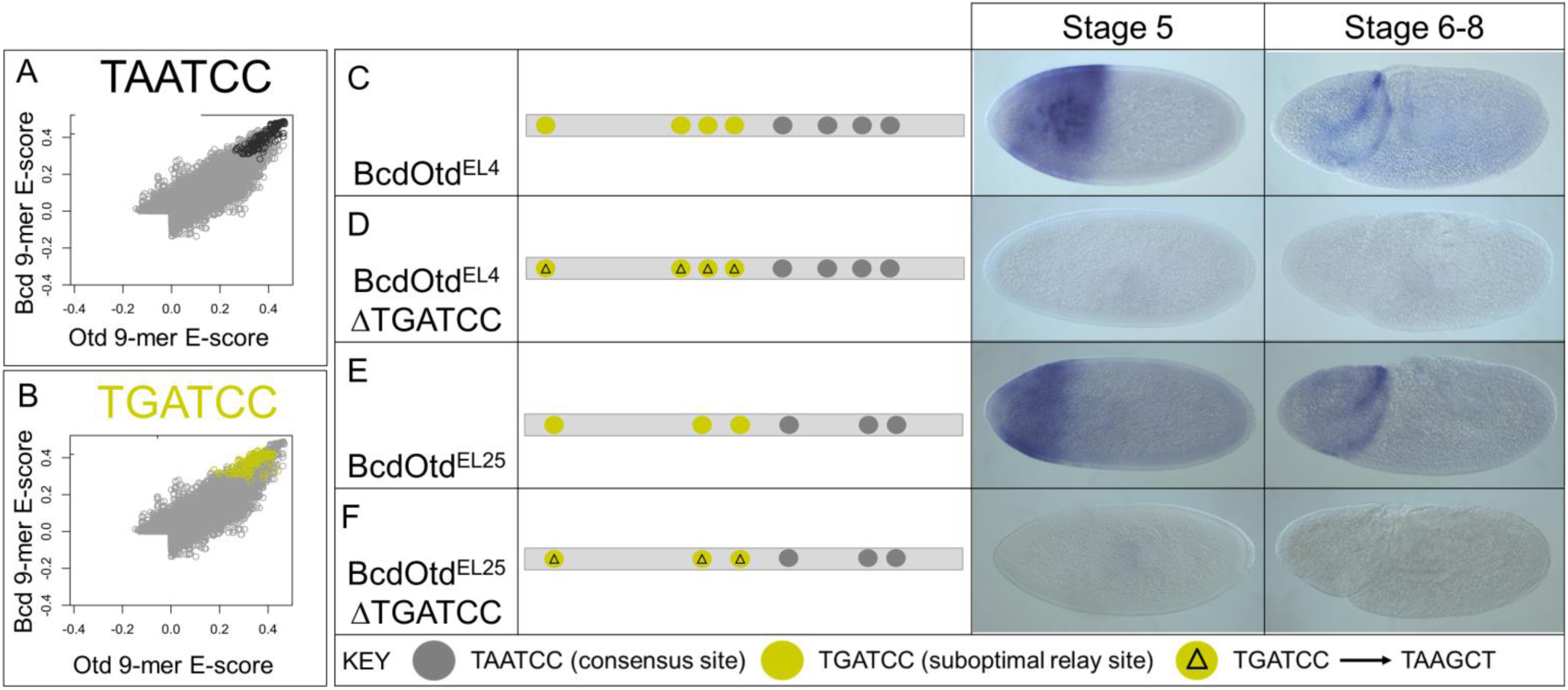
**(A-B) Protein-binding microarray comparisons of DNA-binding preferences between Bcd and Otd.** Scatterplots showing all 9-mers that contain the K50 binding consensus motif TAATCC (black circles in A), and all 9mers containing the variant TGATCC that is enriched in feed-forward enh. ancers (yellow circles in B). **(C-F) Functional tests of the TGATCC motif.** lacZ expression patterns are shown at two time-points for two wild-type enhancers (C, E), and those same enhancers with point mutations in TGATCC motifs (D, F). A binding site key is shown at the bottom.

To test the *in vivo* function of the TGATCC suboptimal site, we mutated it in two different enhancers (BcdOtd^EL4^ and BcdOtd^EL25^). Both are relay enhancers because (a) they are activated at both S5 and S6-8 (Fig. 5C, E), (b) they are bound by both Bcd and Otd, (c) they lose expression in *bcd* mutants, and (d) they lose expression late in *otd* mutants ([16]; Fig. 4). The BcdOtd^EL4^ and BcdOtd^EL25^ enhancers each contain three copies of the TGATCC site, and four and three copies, respectively, of the TAATCC consensus site. We converted the TGATCC sites in these enhancers to TAAGCT– a lower affinity site for both proteins that is equally represented in all ChIP datasets. The consensus sites were not mutated. For both enhancers, these mutations caused a complete loss of expression (Fig. 5D, F), suggesting that this motif is important for both Bcd-mediated activation and Otd-dependent maintenance of expression and that the TGATCC binding site is critical for the Bcd/Otd relay.

### Anterior patterning by the integration of three classes of enhancers

In addition to the relay enhancers identified by overlapping peaks of Bcd and Otd binding, our ChIP-Seq experiments yielded large numbers of unique binding peaks for each protein (Fig. 3). The 630 peaks unique to Bcd included 13 previously known Bcd-dependent enhancers [16]. The expression patterns of 8 of these enhancers perdure after the Bcd gradient decays (Supp. Fig. 5), but they were not bound by Otd, so we hypothesize that other factors must be involved in the maintenance of their patterns (see Discussion).

The ChIP experiments also identified many peaks bound uniquely by Otd at each time-point. We used *lacZ* reporter genes to test 19 of these peaks for enhancer activity. In our first experiments, we tested nine of the 719 fragments bound specifically by Otd only at the late timepoint (Otd^L^; Fig. 6B, Supp. Fig. 5). As expected, all nine enhancers showed anterior expression patterns in S6-8 embryos (Fig. 6B, Supp. Fig. 5). We also tested ten genomic fragments bound uniquely by Otd only at S4 (Otd^E^, 4 fragments), or at both timepoints (Otd^EL^, 6 fragments). Surprisingly, none of the ten tested fragments directed any reporter gene expression at either stage of development. (Fig. 6A, Supp. Fig. 5). These results are consistent with the failure of Otd to rescue *bcd* mutant embryos (Fig. 1), and suggest that Otd-binding to euchromatic regions at S5 does not lead to enhancer activation. The reason for Otd’s failure to activate expression at S5 is not clear, but one possibility is that activation of anterior genes in the early embryo requires prior binding by Bcd. Alternatively, because activation by Bcd requires interactions with cofactors [34-36], Otd’s inability to activate may be caused by a failure to make critical protein-protein contacts.

**Figure 6.**
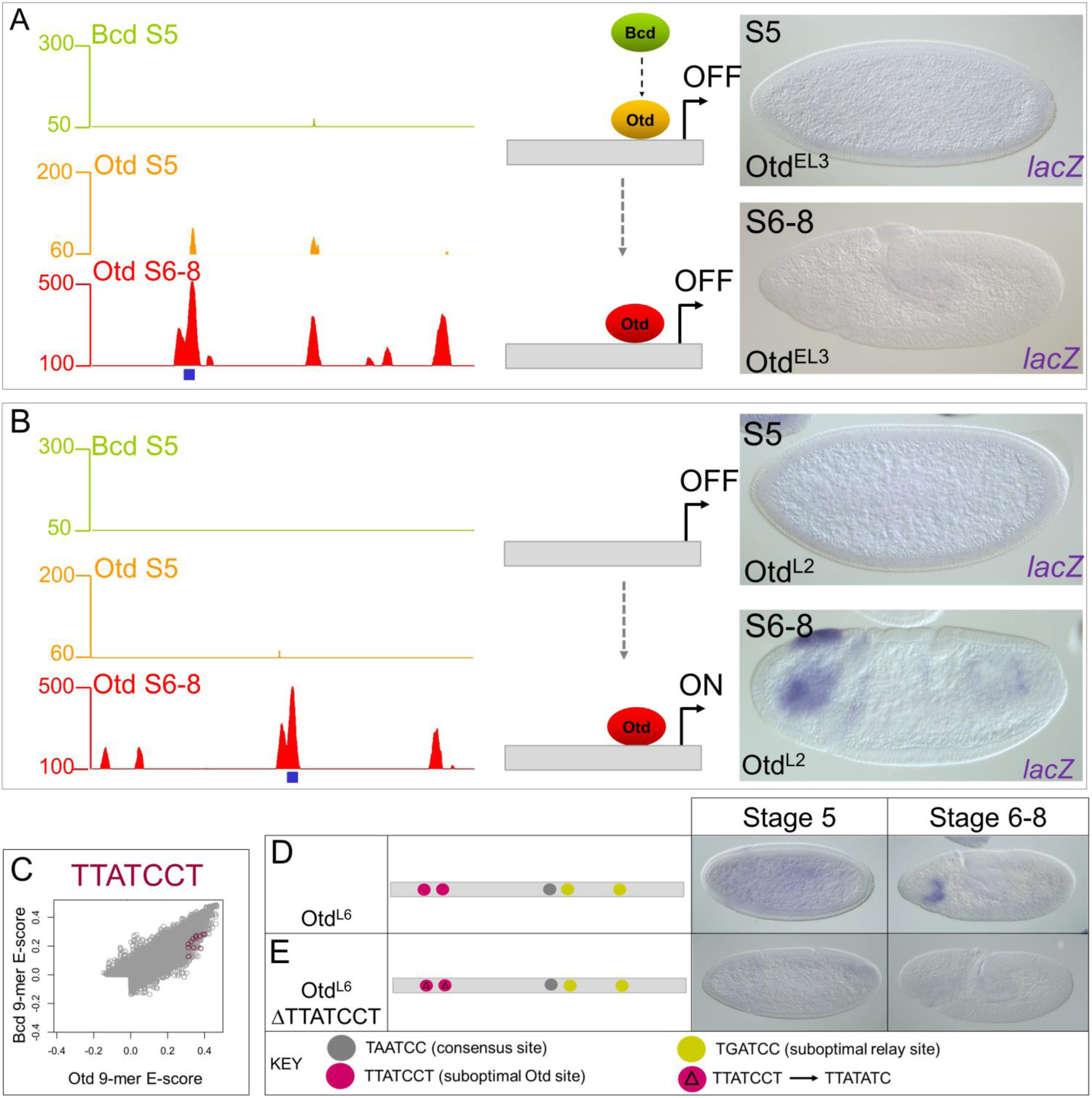
Reporter gene tests of genomic fragments bound only by Otd. **(A)** Otd^EL3^ is bound by Otd at S5 and S6-8, but not by Bcd and is inactive **(B)** The Otd^L2^ enhancer is bound by Otd at S6-8 and is active in the head. **(C)** Otd-preferred sequence TTATCCT that is enriched in the Otd S6-8 ChIP data in preferred in vitro by Otd. **(D-E)** Mutating the Otd suboptimal site TTATCCT abolishes enhancer activity in a late-acting Otd enhancer.

Taken together, our results define three classes of enhancers that mediate Bcd and/or Otd functions *in vivo*. Class 1 enhancers relay transcriptional initiation by Bcd to transcriptional maintenance by Otd, Class 2 enhancers are activated by Bcd, and are maintained by an unknown factor. Class 3 enhancers are activated at a later time-point by an Otd-dependent mechanism that is completely independent of Bcd.

### Otd-dependent activation is mediated by another suboptimal site

We then searched for over-represented motifs within the set of peaks bound specifically by Otd at S6-8. This search identified an enriched K50-variant motif in Otd (TTATCCT) an extended variant of the canonical TAATCC motif optimally preferred by Bcd and Otd. It appears in 26% of Otd S6-8 peaks, 10% of Bcd+Otd peaks, and only 9% of Bcd S5 peaks (p=1.89E-15). An analysis of our PBM data showed that this suboptimal site was preferentially bound by Otd compared to Bcd (Fig. 6C), consistent with its over-representation in Otd-bound genomic fragments.

We tested the role of the TTATCCT motif in an enhancer (Otd^L6^) that is bound by Otd and transcriptionally active only at S6-8 in wild type embryos, and inactive in *otd* mutants (Fig. 6D, Supp. Fig. 5). The Otd^L6^ enhancer contains two exact copies of the TTATCCT motif, (Fig. 6D), one copy of the canonical K50HD-binding motif (TAATCC), and two copies of the TGATCC motif that is over-represented in relay enhancers. We mutated the TTATCCT motifs, leaving the canonical and relay motifs intact, which caused a complete loss of expression (Fig. 6E). Based on our PBM data, ChIP data enrichment, and the enhancer site-change data, we conclude that these suboptimal sites are necessary for activation by Otd at S6-8.

### Timing of enhancer activity is controlled by suboptimal motif preferences and cofactor interactions

Our motif searches identified suboptimal binding sites required for the activities of feed-forward relay enhancers (Fig. 5) and Otd^L^ enhancers (Fig. 6) respectively. These results suggest that subtle binding preferences between the two proteins contribute to correctly timing enhancer activation *in vivo*. Alternatively, it is possible that the differential timing of enhancer activity is controlled by the presence or absence of binding sites for protein-specific cofactors. Previous studies identified two transcription factors, Hunchback (Hb) and Zelda (Zld), as being critical for Bcd-dependent gene activation [35-38]. Consistent with this, a discriminative motif analysis revealed an enrichment of Hb and Zld motifs in regions bound *in vivo* by Bcd and Bcd+Otd compared to Otd only (Fig. 4I).

We hypothesized that early activation of enhancer activity by Bcd is controlled in part by co-factor interactions, and that the addition of Hb and/or Zld sites might convert a late-acting enhancer into a relay enhancer that is expressed earlier. We tested this hypothesis on Otd^L6^, which is bound by Otd at S6-8 and active only at this later stage. This enhancer is not active early, and not bound by Bcd *in vivo* despite having two copies of the relay motif (TGATCC; Fig. 7A). We added 4 high affinity Hb sites to the Otd^L6^ enhancer, which resulted in early activation of expression as an anterior stripe (Fig. 7B). Adding four canonical Zld sites into this element also caused it to be activated early, but in a complex pattern along the length of the embryo (Fig. 7C), and adding both Hb and Zld sites had an additive effect on the enhancer’s activities (Fig. 7D). Otd^L6^ enhancers containing extra Hb and Zld sites were each crossed into *bcd* mutants, which caused a complete loss of expression (data not shown), consistent with the hypothesis that the addition of Hb and Zld sites successfully converted the Otd^L6^ enhancer into an early acting Bcd-dependent enhancer.

**Figure 7.**
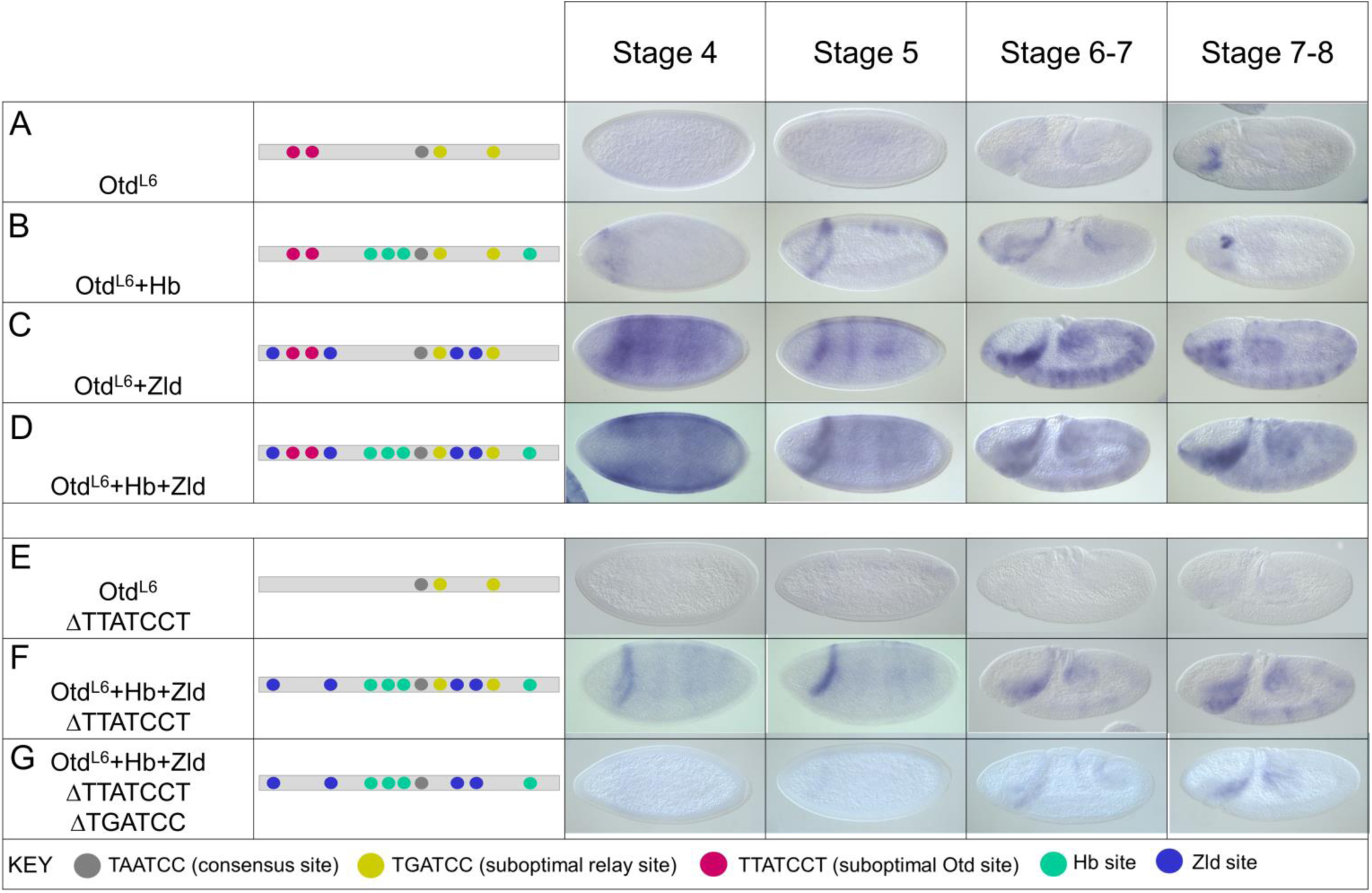
Binding site manipulations that change the timing of enhancer activity. **(A-D)** *lacZ* RNA expression patterns at four different time-points driven by the wild type Otd^L6^ enhancer (A), and the Otd^L6^ enhancer with four added Hb sites (B), four added Zld sites (C), and four extra Hb sites plus four extra Zld sites (D). **(E)** *lacZ* expression driven by the Otd^L6^ enhancer carrying mutations in three suboptimal Otd (TTATCCT) sites. Compare to the patterns in A. **(F)** *lacZ* expression driven by the Otd^L6^ enhancer carrying mutations in three TTATCCT sites, and four additional sites each for Zld and Hb. **(G)** *lacZ* expression driven by the Otd^L6^ enhancer carrying mutations in three TTATCCT sites, two TGATCC sites, and four additional sites each for Zld and Hb. See text for a description of results and interpretations.

Our experiments above suggest that the specific sequence variants TGATCC and TTATCCT are required for the activities of relay and Otd-activated enhancers, respectively (Fig. 3D, H, Fig, 5C, Fig. 6C). The Otd^L6^ enhancer contains both sequence variants, and thus provides an opportunity to test how each motif might interact with the Zld and Hb cofactors. As shown in Figure 6D, mutating the TTATCCT motifs in the Otd^L6^ enhancer caused a loss of expression, even though there are two intact copies of the TGATCC sequence. We hypothesized that adding Hb and Zld sites to this inactivated enhancer might “rescue” regulatory activity. Adding Hb and Zld sites to Otd^L6^ΔTTATCCT resulted in a stripe of expression that was expressed early and maintained in S6-8 (Fig. 7F), a temporal pattern similar to that observed for other relay enhancers. This result suggests that the combination of only two TGATCC sites can mediate early activation if augmented with binding sites for Zld and Hb. A further mutation of the two TGATCC sites in the Otd^L6^+Hb+Zld ΔTTATCCT enhancer resulted in a loss of expression, indicating the importance of those sites for early activation, despite the presence of strong binding sites for Zld and Hb (Fig. 7G).

Taken together, our experiments suggest that the specific responses of enhancers to Bcd-and Otd-mediated activation are controlled in part by suboptimal motifs preferred by each protein, and that early activation by Bcd requires interactions with the cofactors Hb and Zld.

## Discussion

In this paper, we compared the *in vivo* functions of two K50 HD proteins (Bcd and Otd) within a framework of evolution. We showed that Otd and Bcd have evolved independent functions *in vivo*, and HD swaps between the two proteins indicated that the major structural differences mediating their distinct *in vivo* activities can be traced to their HDs. The two proteins each bind to unique enhancers, and Otd also binds enhancers previously bound by Bcd via a feed-forward relay (FFR) mechanism that extends the timing of the gene expression patterns they regulate. We presented evidence that Otd binding does not lead to enhancer activation in the early embryo, but it is an effective activator after the first wave of zygotic target genes are activated. Finally, we showed that enhancers respond in specific ways to Bcd and Otd through suboptimal binding sites for K50 HD protein and binding sites for cofactors.

### The Bcd-Otd FFR in *Drosophila*

Bcd has evolved rapidly to become a powerful morphogen in *Drosophila*, but is not well-conserved, even within Diptera [23]. In *Drosophila*, *otd* has evolved to become a zygotic target gene of Bcd[18]. As such, *otd* RNA appears at S4 and its protein is detectable at early S5, at the same time as most other Bcd target genes. Despite having a lethal mutant phenotype, there is little known about Otd’s molecular functions in *Drosophila* embryogenesis. Otd binds to many enhancers that were initially activated by Bcd at S4 (Fig. 3). The reduced expression driven by these enhancers in *otd* mutants at S5 suggests that Otd protein binding is functional, and the loss of expression at Stages 6-8 demonstrates that Otd is required for maintaining their activity (Fig.4). These results suggest strongly that Bcd and Otd participate in a relay, in which Bcd initiates transcription activation of target genes including *otd*, and Otd maintains its own expression and the expression of other target genes after the Bcd gradient decays. This relay is similar to the C1 feed-forward loop (FFL) with an OR-like input function of Alon [39], but it is distinct in two respects. First, all target enhancers regulated by the Bcd-Otd FFR are initially activated by Bcd before Otd is present. Second, the relay involves the sequential binding of Bcd and Otd to the same binding motifs in the same target gene enhancers.

Because Bcd and Otd recognize similar sequence motifs *in vitro*, and they are involved in an FFR in *Drosophila*, one would predict that a maternal gradient of Otd (Mat:Otd) would activate many Bcd-dependent target genes when Bcd is genetically removed. To the contrary, we found that Mat:Otd cannot activate the great majority (more than 90%) of the tested Bcd-dependent transcriptional targets, including many that are regulated by the Bcd-Otd feed-forward loop. Also, our ChIP-Seq experiments identified hundreds of peaks that bind Otd but not Bcd in early embryos, but most of these peaks map to heterochromatic or uncharacterized regions of the genome. We tested ten such fragments from euchromatic regions, but none showed any enhancer-like activity. Taken together, these results suggest that Otd is unable to efficiently activate transcription on its own in early *Drosophila* embryos, and that its role in early embryogenesis is inextricably linked to Bcd.

The most likely explanation for Otd’s inability to replace or function well without Bcd is that it is unable to interact efficiently with the Bcd cofactors Zld or Hb. Bcd activates the very first zygotic target genes as part of the maternal to zygotic transition (MZT), and a key factor in the MZT is Zld, which is thought to act as a pioneer that opens chromatin [40, 41]. We have previously shown that Zld facilitates Bcd binding to target gene enhancers [36]. Similarly, most Bcd target genes require the activity of Hb to increase target gene sensitivity to Bcd-dependent activation [35, 38]. Interestingly, because insertion of the Bcd HD into the Otd protein confers on it many of Bcd’s normal functions, these critical interactions may involve direct contacts with peptide motifs within Bcd’s HD. We propose that similar motifs are not present in Otd’s HD, which accounts for its inactivity at the early time-point.

### Mechanisms controlling different enhancer activities

The ChIP-Seq experiments enabled us to group genomic fragments based on their ability to bind Bcd and/or Otd and their temporal expression patterns (Fig. 8). Fragments in the first group bind Bcd early and Otd late, and mediate the feed-forward loop described above. 53 of the 66 known Bcd-dependent enhancers belong to this class. Fragments in the second group are bound early by Bcd and never bound by Otd. Only 13 of the 66 known Bcd-dependent enhancers are in this class. These enhancers are also active after the Bcd gradient degrades, but they do not bind Otd, and it is not clear how their expression is maintained. Fragments in the third group (Otd^L^) are never bound by Bcd, but are bound by Otd at S6-8. We tested nine Otd^L^ candidate enhancers, and all showed enhancer activity.

**Figure 8.**
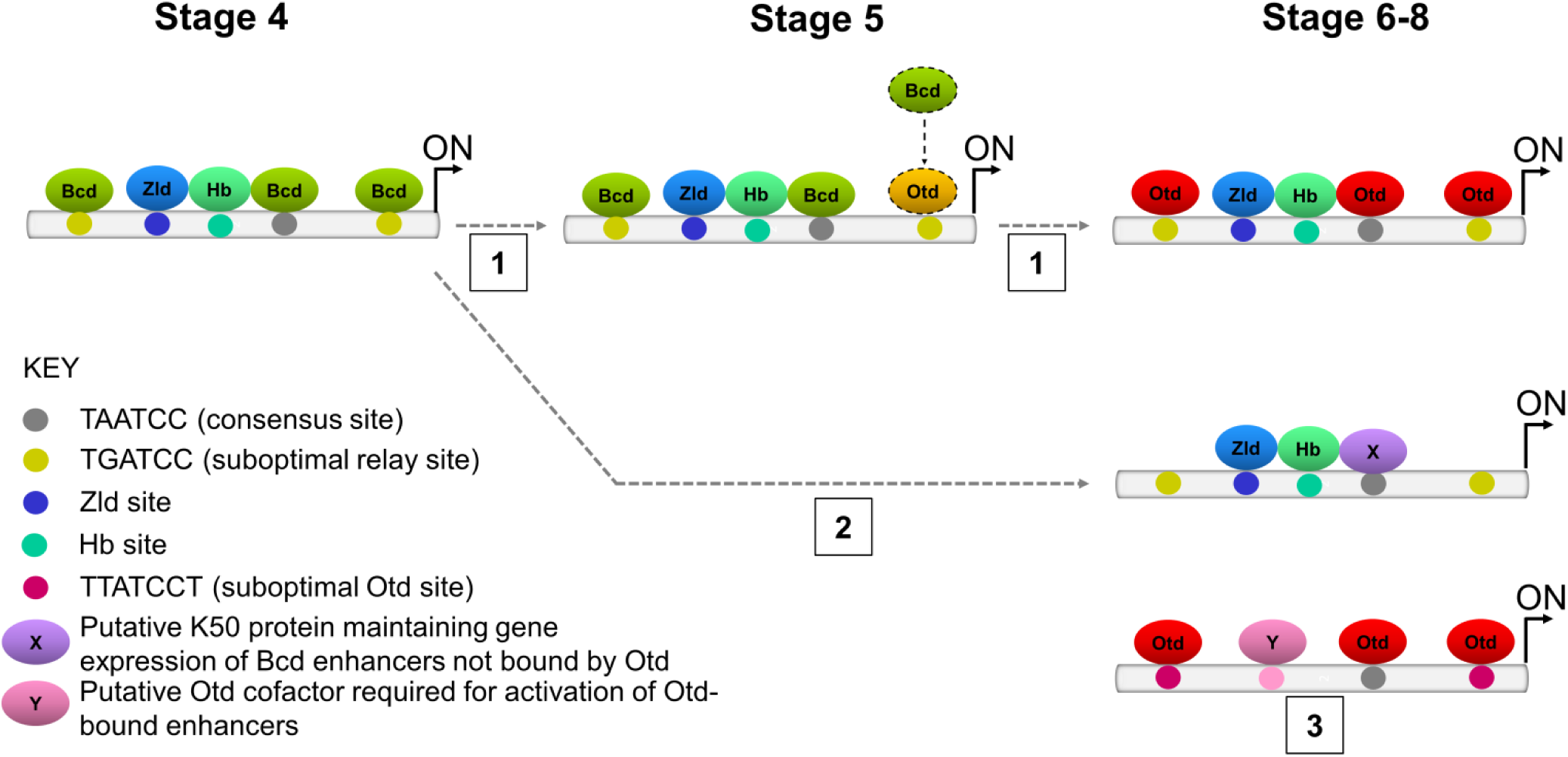
Three classes of Bcd- and Otd-dependent enhancers. Class 1: Bcd-Otd feed-forward relay enhancers are activated at Stage 4 by Bcd with the cofactors Hb and Zld. Bcd protein binds to consensus (TAATCC) and suboptimal (TGATCC) sites required for gene activity. At Stage 5 Otd protein binds to suboptimal sites, replacing Bcd. At Stage 6-8 Otd protein binds to all consensus and suboptimal sites to maintain gene activity. **Class 2:** These enhancers are activated by Bcd, and maintained after the Bcd gradient decays, but they are never bound by Otd. These enhancers may be regulated by a feed-forward loop involving an unknown factor (X). **Class 3:** Other Otd activates gene targets in the late stage through binding to both consensus and suboptimal (TTATCCT) sites. Otd enhancer activation at S6-8 might require another cofactor (Y).

We have begun to understand the molecular mechanisms that distinguish the activities of feed-forward enhancers from those activated later. Motif searches identified two over-represented variants of the TAATCC consensus, TGATCC and TTATCCT in feed-forward enhancers and Otd^L^ enhancers respectively; Fig. 5C, D). PBM experiments showed that both variants are suboptimal binding sites compared to TAATCC, but Bcd prefers to bind the TGATCC motif, while Otd prefers TTATCCT. Mutational analyses showed that these motifs are required for the activation of their respective enhancers, which are consistent with recent studies of lower affinity but still functional sites in many systems [42-46]. However, it is important to point out that many ChIP-Seq peaks specifically bound by Otd or Bcd do not contain the expected “preferred” variant motifs, and that the occurrence or absence of these sequence variants is not predictive of enhancer activity per se. Other sequence variants of the consensus are over-represented in feed-forward and Otd^L^ enhancer groups (data not shown). These results suggest that global binding preferences may be controlled by the aggregate activities of multiple sequence variants.

Sequence motifs for Zld and Hb were also over-represented in ChIP-Seq peaks bound by Bcd (Fig. 3), consistent with the combinatorial activation model mentioned above. Adding Zld and Hb binding sites to an enhancer normally activated later by Otd results in earlier activation and conversion of that enhancer into a Bcd-dependent enhancer (Fig. 7B-D; F). We hypothesize that Otd^L^ enhancers contain binding sites for unknown Otd-specific cofactors, and identifying these sites and the proteins that bind them will be the focus of future studies. Taken together, our results indicate that both intrinsic DNA-binding preferences and interactions with cofactors control the distinct temporal and spatial patterns of expression driven by individual enhancers *in vivo*.

### Convergent evolution and a robust core anterior patterning network

*bcd* and its paralog *zen* arose through duplication of an ancestral maternally expressed *Hox3*-like gene [23, 47]. In insects lacking Bcd, Otd has Bcd-like properties (maternal expression, anterior mRNA localization and patterning) [24, 25]. Since the two proteins do not share common ancestry (Otd is a Paired class homeoprotein, which diverged from the Hox cluster over 800 MY ago), some functional convergence of these proteins must have occurred during insect evolution.

The evolution of Bcd likely involved the retention of the maternal promoter, and the acquisition of three characteristics required for anterior embryonic patterning: 1. UTR sequences that control anterior mRNA localization, 2. protein motifs that mediate translational repression, and 3. amino acid substitutions that alter its DNA-binding preferences, namely a Q50 to K50 mutation in its HD. Since *otd* is zygotic in *Drosophila*, some of these characteristics described above must have been lost in *otd* in the lineage that led to *Drosophila*. Such dramatic changes in these two genes may be attributed to reduced selective pressure on maternal genes [48, 49], which permits the exploration of the evolutionary landscape and the acquisition of new functional roles.

What is striking is that Otd binds to at least half of the Bcd-bound target genes after they are activated by Bcd. Perhaps this set of target genes represents an ancestral core network that is well conserved in evolution. Many feed-forward enhancers are associated with the gap genes, which cross-regulate each other via repressive interactions. These enhancers all contain DNA motifs that are recognized by a K50HD transcription factor like Bcd or Otd. Thus it is possible that the cis-regulatory motifs (and consequently the enhancers) are functionally robust in the evolution of anterior embryo patterning, while trans-acting factors can accumulate mutations. This allows for a conserved set of targets that make up a canalized anterior patterning network that allow the regulating transcription factors to evolve.

## Methods

### Drosophila melanogaster stocks

The following stocks were used in these experiments: yw (wild type), ±/*Cyo bcd*^+^;*bcd*^*E1*^/*bcd*^*E1*^, *Cyo bcd*^+^/*Sco*;*bcd*^*E1*^/*bcd*^*E1*^, and ΦC31 (y+);38F1 (w+).

### Maternal gene chimeras

The *bcd* and *otd* coding regions were amplified by PCR from pBS-SK+ cDNA clones. We cloned an injection plasmid (piattB40-Bcd) using traditional techniques from pBSSK+ (Asp718/SacI) and industrially synthesized oligonucleotides. This plasmid contains inverted phiC31-specific recombination sequences (AgeI/HindIII), the fluorescent green eye marker Gmr-Gfp (HindIII/AscI), and a polylinker flanked by the *bcd* promoter and 3’UTR (AgeI/AscI). The amplification product was digested with RsrII and AscI and ligated into the plasmid fragment. Homeodomain swaps and residue changes were generated using standard cloning techniques and nested PCR. All transgenic lines were generated using the ΦC31 integration system, and constructs were integrated into the 38F1 landing site on the third chromosome [31].

### Embryo collections, cuticle preparations, and immunohistochemistry

Embryos were collected 2-3hr and 3-5 after egg laying (AEL). Embryos were dechorionated for 2 minutes in bleach, and a 2:1 mixture of methanol and heptane was used to remove the vitelline membrane. ISH and FISH were performed as previously described [50] using digoxigen-, fluorescein-, and biotin-labeled probes. Cuticle preparations were performed on embryos aged 20-24 hours. For cuticle preps larvae were fixed overnight at 65°C in a 1:4 mixture of glycerol and acetic acid, and mounted in a 1:1 mixture of Hoyer’s medium and lactic acid. Rabbit anti-Bcd (1:400) and Guinea pig anti-Otd (1:1000) were used for immunostaining (with rabbit-FITC and 647-guinea pig as secondary antibodies, 1:400 each, Invitrogen). Guinea pig anti-Cad (1:400) and Alexa Flour® conjugated 488 donkey anti-guinea pig (1:500, Invitrogen) were used to examine Cad protein expression. All antibodies were diluted in PBT. Data for immunostaining images were collected on a Leica TCS SP5 confocal microscope using the Leica confocal analysis software. Gradient quantifications were performed as previously described [51].

### ChIP-sequencing

Chromatin immunoprecipation was performed using Bcd antibody (rabbit) and two Otd antibodies (guinea pig; GP-5 and GP-6, both provided by Tiffany Cook) on two biological replicates (two technical replicates per sample) of wild-type chromatin collected at 2-3hr and 3-5 AEL. Antibodies were Protein A purified using the Protein A Antibody Purification Kit (Sigma). The embryos from each collection were DAPI stained to confirm embryonic stages and 85% of the embryos were the correct age in each collection. Embryos were treated and fixed in 1.8% formaldehyde as previously described [36, 52]. Samples were sonicated for 10 min at Setting 3 (30s on, 30s off) followed by 2.5 min at Setting 4 (30s on, 30s off) using a Sonic Dismembrator Model 550. 200ul of embryonic lysate was used for each immunoprecipitation reaction. 240ul buffer FA+PI (50 mM HEPES/KOH pH 7.5, 1 mM EDTA, 1% Triton X-100, 0.1 % sodium deoxycholate, 150 mM NaCl, protease inhibitor) was added to each sample to bring the total volume to 440ul. 5% of each extract was removed as input. 10ul of each antibody was added to the sample (minus input) and incubated overnight at 4°C. 40 ul of protein A sepharose bead slurry (Amersham Biosciences) was added per ChIP sample with 1 ml FA buffer. The sample was centrifuged at 2500 g for 1 min and supernatant was discarded. 1 ml FA buffer was added to each tube. Beads were suspended by inverting the tubes a few times, then centrifuged again. This wash was repeated three times. After the final wash, the beads suspended in 40ul FA buffer. The ChIP sample was added to the bead slurry and rotated at 4°C for 2 hours. 2ul of 20mg/ml RNaseA was added to the inputs. The beads were washed at room temperature by adding 1 ml of each of the following buffers and incubating on a nutator (or a rotator): FA (2X for 5 min), FA-1M NaCL (1X for 5 min; 50 mM HEPES/KOH pH 7.5, 1 mM EDTA, 1% Triton X-100, 0.1 % sodium deoxycholate, 1 M NaCl), FA-500mM NaCl (1X for 10 min; 50 mM HEPES/KOH pH 7.5, 1 mM EDTA, 1% Triton X-100, 0.1 % sodium deoxycholate, 500 mM NaCl); TEL (1X for 10 min; 0.25 M LiCl, 1% NP-40, 1% sodium deoxycholate, 1 mM EDTA, 10 mM Tris-HCl, pH 8.0); Tris-EDTA (2X for 5 min). The beads were collected after each wash by centrifugation for 1 minute at 2500 g and removing supernatant. To elute the immunocomplexes, we added 125ul ChIP Elution Buffer and placed the tube in a 65 °C heat block for 15 min. We spun down the beads at 6000 g for 1 min and transferred the supernatant to a new tube. The elution was repeated and supernatants were combined.

### ChIP-seq library preparation and data processing

NEXTflex ChIP-seq kits from BIOO Scientific (#5143-01) were used to prepare libraries, which were then barcoded using NEXTflex ChIP-seq barcodes (BIOO Scientific, #514120). We performed single-end HiSeq2000 sequencing (Illumina) at the New York University Genome Center. Sequencing reads were mapped using Bowtie to the *Drosophila melanogaster* genome release 5.3, and duplicate reads were removed after combining data from each biological replicate. The MACS program was used to call peaks over input [53, 54]. We removed all heterochromatic and uncharacterized chromatin from the analysis, and further narrowed down relevant peaks by using only those that appeared in both antibodies (for Otd), and across all replicates. All ChIP-seq data processing was done on the Galaxy Cistrome platform. An FDR cutoff of 5% was used for all further analysis on ChIP-seq peaks.

### Motif enrichment analyses

Matlab was used for all motif analyses, unless stated otherwise. Bcd and Otd peaks were compared and peaks were considered shared if they overlapped by at least 200 base pairs. We compared all of our ChIP-seq datasets to prior DNase I, Hb ChIP-chip, and Zld ChIP-seq experiments using the same overlapping criterion. We used MEME-ChIP [55] in the MEME suite [56] to search for overrepresented motifs in each ChIP dataset. We then used a discriminative *de novo* motif search to look for overrepresented 6-mers, 7-mers, 8-mers, and 9-mers in ChIP-seq peaks. Cofactor PWMs used to search Bcd and Otd bound genomic regions were derived as follows: Bcd and Otd (from PBM done in this paper), Zld and Hb (Fly Factor Survey, http://mccb.umassmed.edu/ffs/).

### GST-tagged proteins

Homeodomain coding sequences (as well as 15 amino acid flanking regions) for Bcd and Otd were PCR amplified and cloned into a N-terminal GST fusion Gateway expression vector (pDEST15, Invitrogen) and the correct clones were confirmed by sequencing. Rosetta™(DE3) competent cells (BL21-derivatives from Novagen) were transformed with the GST-BcdHD or GST-OtdHD plasmids. For each transformation, a single colony was used to inoculate 0.5-L of LB+Amp^100^ and shaken at 200 rpm at 37°C until the OD_600_ reached 0.5-0.6. Proteins were induced at 37°C for 3 hours by the addition of 0.5 mM IPTG. After induction, bacteria were harvested by centrifugation (15 min at 5K xg) and the cell pellets were stored at −80°C. Each frozen pellet was resuspended in 40 mL of lysis buffer (150 mM NaCl, 10 mM HEPES pH 7.9, 5 mM DTT, 10% glycerol, 0.5% Triton X-100, 0.5 mg/ml lysozyme, 1mM MgCl2, 1 mM Benzamidine, 3 µM Pepstatin A, 2 mM Leupeptin, 1 mM PMSF) and sonicated using a Branson Sonifier 450 homogenizer equipped with a midi-tip. The sonication parameters were: 6-8 strokes of 1 min with an output of 6, duty cycle of 50%, and a resting time of 2 min on ice between strokes. The crude lysate was centrifuged at 27K xg for 30 min at 4°C and the supernatant was transferred to a new tube containing 0.5 mL of settled, pre-washed and pre-equilibrated, glutathione sepharose 4B beads (GE Healthcare 17075601) and rotated for 6 hours. Elutions were performed with lysis buffer containing 5 mM (E1 and E2), 7.5 mM (E3), 10 mM (E4 and E5), 20 mM (E6 and E7) and 50 mM (E8 and E9) imidazole. A total of 10 mL of the peak eluates were dialyzed twice for 12 hours each time against 1 L of storage buffer (150 mM NaCl, 5 mM HEPES pH 7.9, 1 mM DTT, 10% glycerol, 1 mM PMSF). After dialysis the samples were centrifuged 5 min at 21K xg, aliquoted, flash-frozen in liquid nitrogen and stored at −80°C.

### Protein binding microarrays (PBMs)

PBMs were performed as described previously [57] using a custom-designed ‘all 10-mer’ universal array [58] in the 8x60K array format (Agilent Technologies, Inc.; AMADID # 030236; [59]. Both proteins were tested at a concentration of 87 nM. Duplicate PBMs were performed for each protein. The array data were quantified as described previously [57] and 8-mer data were averaged over duplicate PBM experiments using the Universal PBM Analysis Suite [60]. Motifs were derived by the Seed-and-Wobble algorithm [57, 60], modified to use 90% of the foreground and background features [33]. As described previously [33], each 9-mer was then assigned the E-score of the lower scoring of its two constituent 8-mers.n previous assays using PBM data for other *Drosophila* homeodomains, 9-mers with an E-score greater than 0.31 were used with confidence to predict real binding events [33]. 9-mer sequences bound preferentially by Bcd versus Otd were identified as described previously [33].

### Generation of *otd* HD deletion by CRISPR

The CRISPR flies were generated as described [61, 62]. gRNAs were inserted into the pU6-BbsI-chiRNA vector (gRNA1 CTTCGAAAAAAAAAAACGAGTTAGC, gRNA2 CTTCGCATATATAAACATTATGTAC). Two homology arms flanking the cleavage sites were inserted into the multiple cloning sites of the pHD-DsRed-attP vector as donor template. The mixture of the donor vector (500ng/μl) and two gRNA vectors (100ng/μl each) were injected into embryos of y[1] M(vas-Cas9).RFP-ZH-2A w[1118]/FM7a. The F1 flies were crossed to YW flies and F2s were screened for the dsRed+ transformants.

### Synthetic enhancer constructs, transgenesis, and K50-dependent enhancer analysis

All genomic fragments were cloned into the piB-HC-lacZ vector previously described [16] using BglII and Asc1. All constructs were inserted using ΦC31 integrase-mediated cassette exchange at 38F1 on chromosome II. We converted low-affinity Zld and Hb sites into high affinity sites in the OtdL^A^ enhancer. Synthetic enhancer fragments for BcdOtd^EL4^ (HC_01), BcdOtd^EL25^ (HC_49), and OtdL^6^ were synthesized by Integrated DNA Technologies. OtdEL and OtdL enhancers were cloned from genomic DNA. Dependency of enhancers on Otd, Bcd, and Mat>Otd was determined by crossing the enhancers to *otd* and *bcd* mutants and Mat>Otd virgin females, collecting progeny embryos and checking to see how *lacZ* expression was affected by ISH.

## Author contributions

Conceptualization, R.R.D., G.Y., R.J., and S.S.; Methodology, R.R.D. P.S., M.L.B., and Z.X.; Software, R.R.D, J.L., I.B., and J.K.; Formal analysis, R.R.D, J.L., and J.K.; Investigation, R.R.D., J.M., X.R., T.B., J.K., and R.L.; Writing – original draft, R.R.D and S.S.; Writing – Review & Editing, R.R.D, S.S., and R.J.; Visualization, R.R.D and D.Y.; Supervision, R.R.D., S.S., and M.L.B.; Project administration, R.R.D., R.J., and S.S.; Funding acquisition, S.S., M.L.B., and R.J.

## Acknowledgements

We are grateful to Tiffany Cook for the Otd antibody, and the Bloomington Stock Center for fly stocks. We thank the NYU Center for Genomics and Systems Biology for all of their help and support with next generation sequencing. We thank Jinshuai Cao for technical assistance and Steve Gisselbrecht for help with PBM K-mer analysis. We also thank Lila Shokri and Anastasia Vendenko for help with the initial stages of PBM analyses. We are grateful to Claude Desplan and Lionel Christiaen for invaluable feedback on the manuscript. M.L.B was supported by NIH RO1 HG005287. R.L. was supported by an NYU Dean’s Undergraduate Research Fellowship. S.S. and R.J. were supported by NIH GM 106090.

---

**Supplemental Figure 1.**
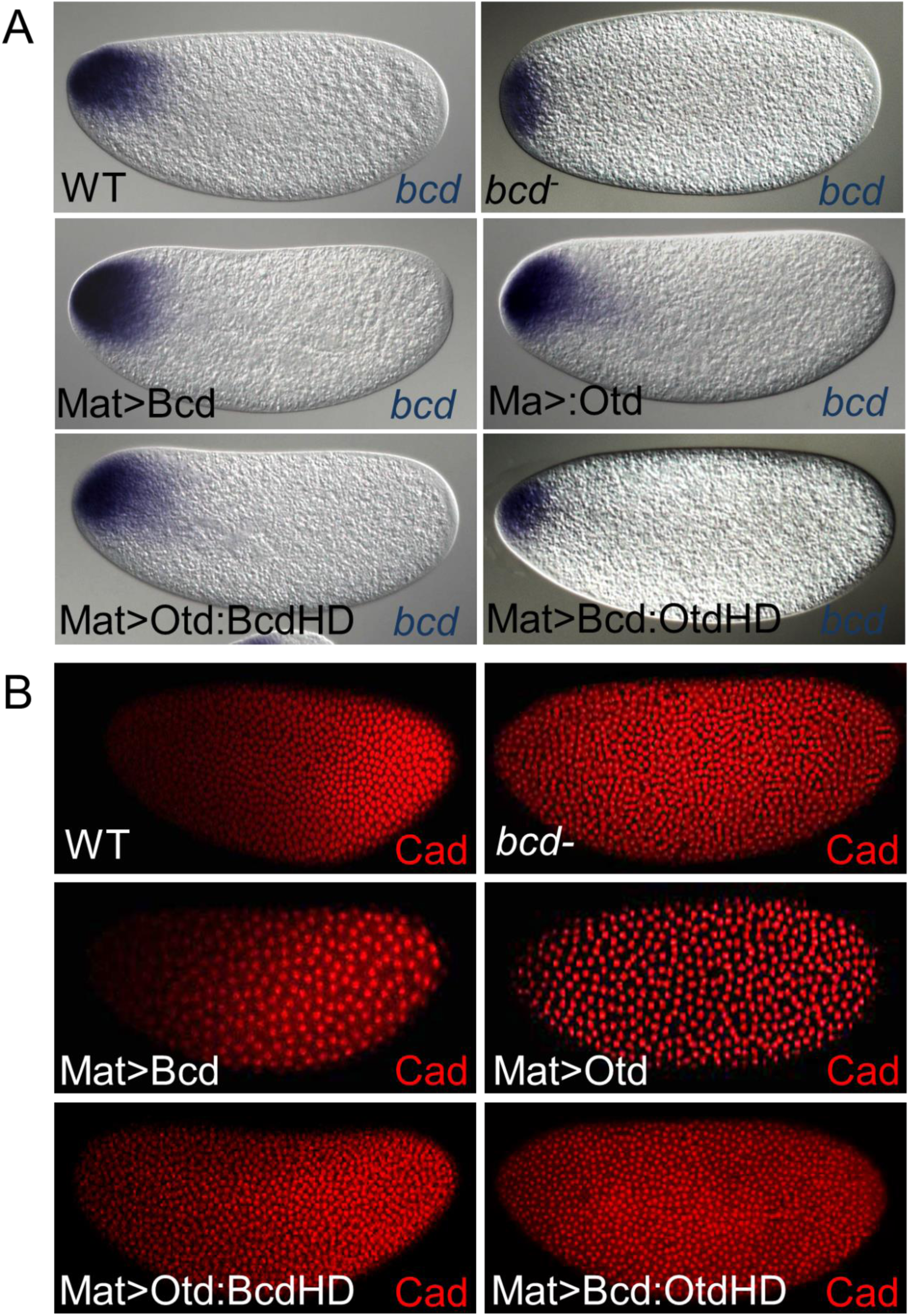
(A) Maternal RNA expression. RNA is localized to the anterior tip for all constructs tested. Construct name and RNA detected are in the bottom left corner. (B) Expression of Cad in wild type, *bcd*^-^, and embryos containing maternal transgenes. Genotypes are indicated in the bottom left corner of the panel.

**Supplemental Figure 2.**
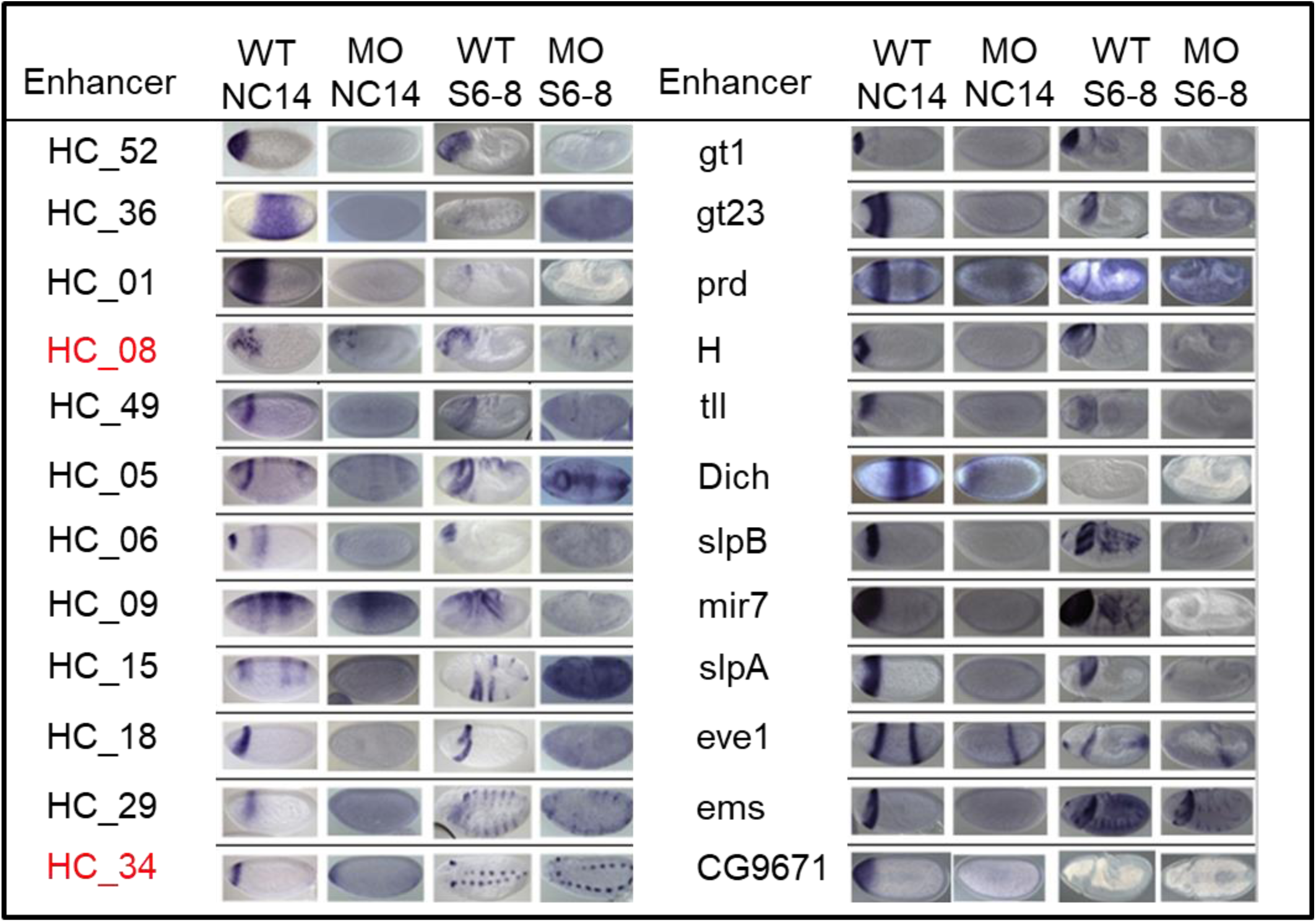
The 24 Bcd-dependent enhancers tested in the Mat>Otd (MO) genetic background. The two enhancers activated by Mat>Otd are shown in red.

**Supplemental Figure 3.**
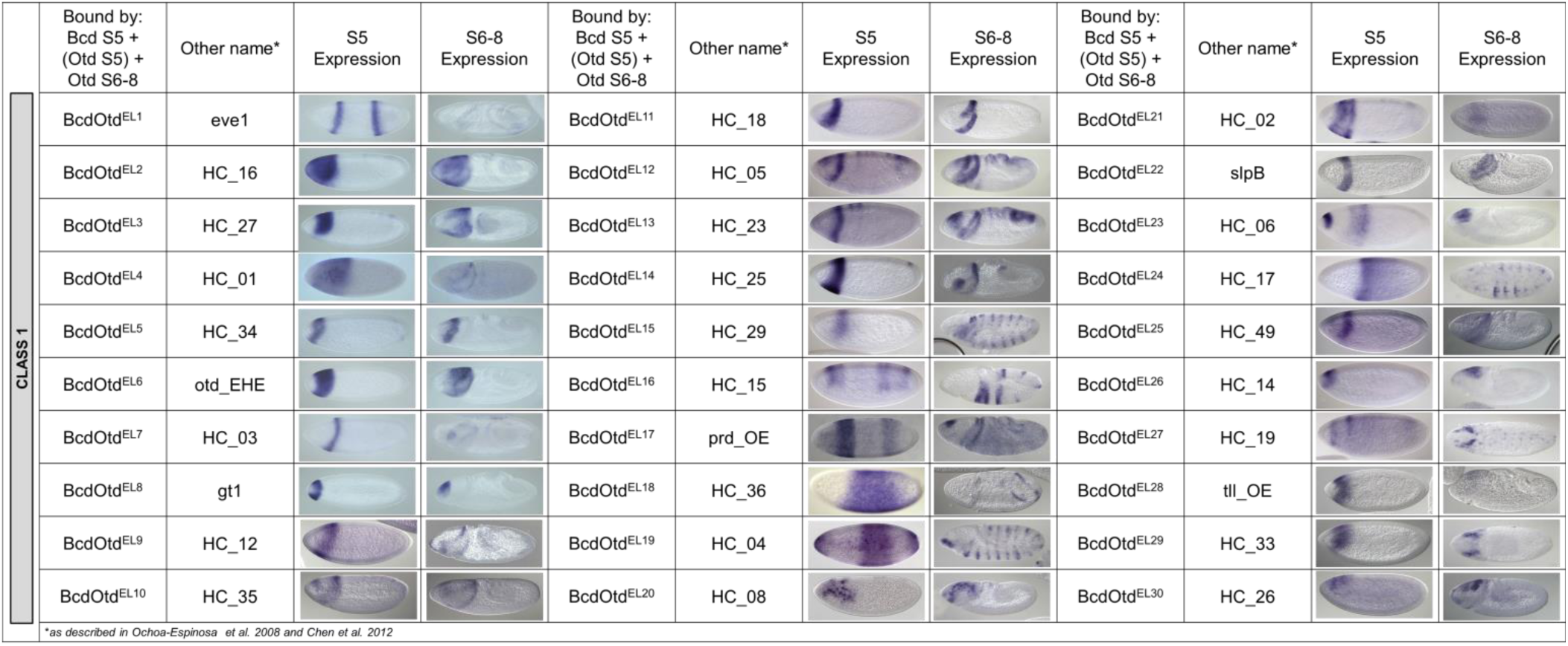
*lacZ* expression patterns of a subset of “relay” enhancers (Class 1).

**Supplemental Figure 4.**
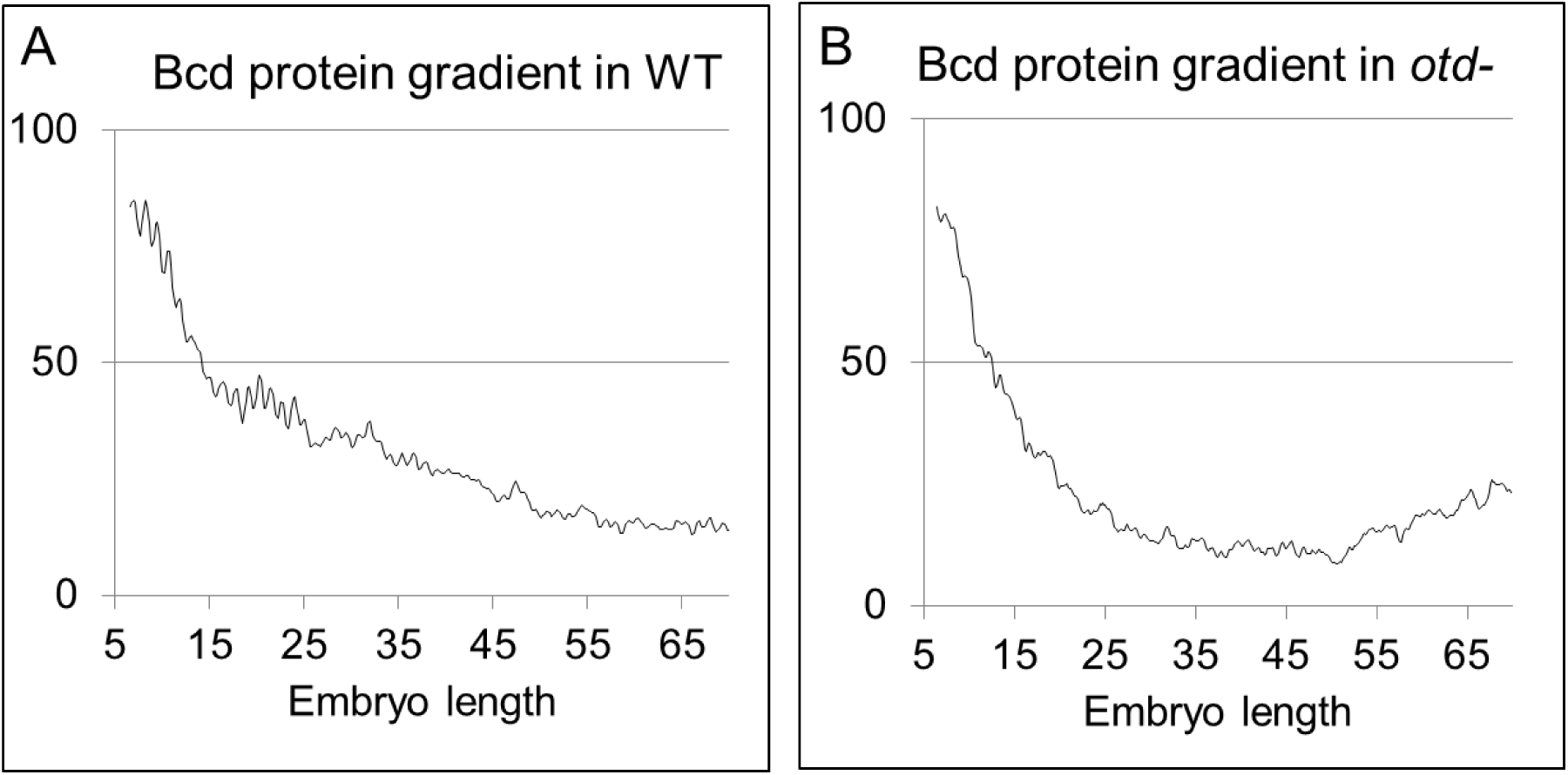
**(A)** Bcd protein gradient in WT and **(B)** otd-embryos.

**Supplemental Figure S5.**
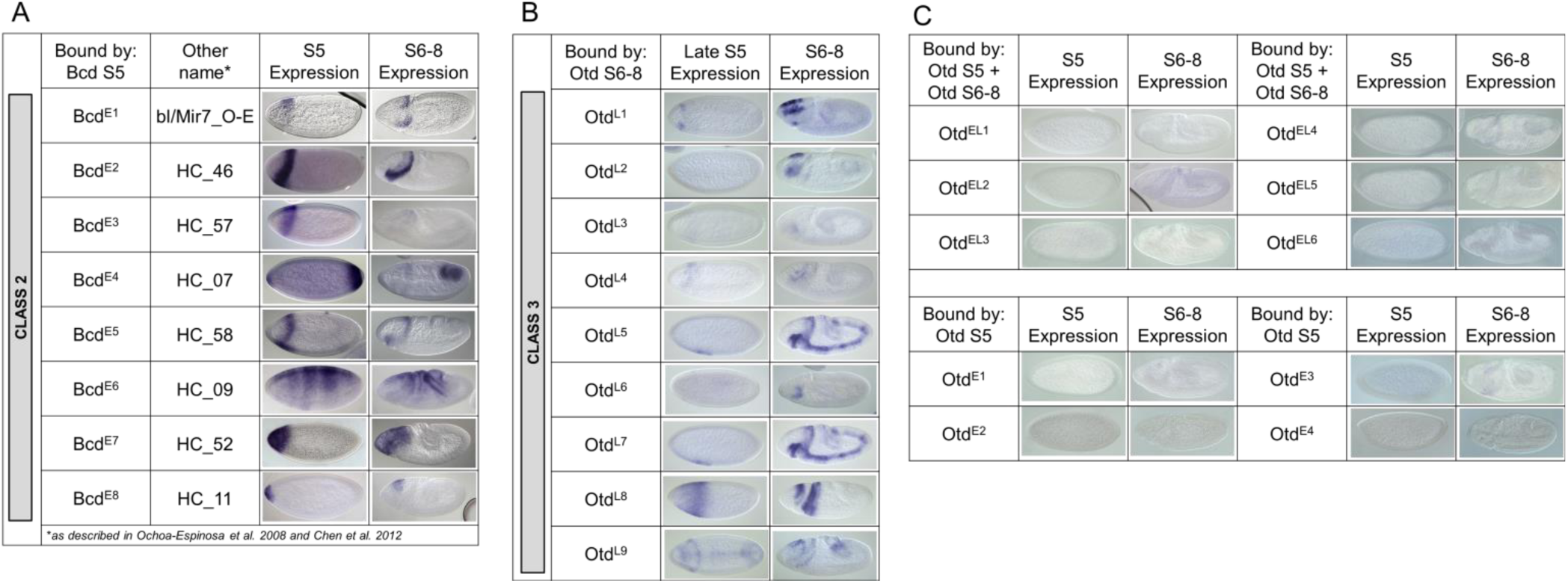
*lacZ* expression patterns of Bcd and Otd bound enhancers. (A) Class 2 enhancers are bound by Bcd and expressed at both timepoints. (B) Class 3 enhancers are bound by Bcd and Otd and are expressed at both timepoints. (C) Otd-bound fragments that are not bound by Bcd do not show any expression.

